# Phosphodiesterase-5 inhibition inhibits epithelial ATP release and restores detrusor contractility in rats with type 2 diabetes *via* an increase in bladder blood flow

**DOI:** 10.1101/2024.03.26.586851

**Authors:** Takafumi Kabuto, So Inamura, Hisato Kobayashi, Xinmin Zha, Keiko Nagase, Minekatsu Taga, Masaya Seki, Nobuki Tanaka, Yoshinaga Okumura, Osamu Yokoyama, Naoki Terada

**Affiliations:** Department of Urology, Faculty of Medical Science, University of Fukui, Fukui, Japan

**Keywords:** type 2 diabetes, rat, bladder, cytokine, tadalafil

## Abstract

**Purpose:** The bladder dysfunction associated with type 2 diabetes mellitus (T2DM) involves urine storage and voiding disorders. We evaluated the pathologic conditions of bladder wall in a rat model of T2DM and evaluated the effects of the phosphodiesterase-5 (PDE-5) inhibitor tadalafil (TA).

**Materials and Methods:** Male Otsuka Long-Evans Tokushima Fatty (OLETF) rats and Long-Evans Tokushima Otsuka (LETO) rats comprised T2DM and control groups. TA was orally administered for 12 weeks. The bladder blood flow and ATP released from the bladder epithelium were measured using laser speckle imaging and an organ bath bladder distention test. The expression levels of markers of hypoxia, pro-inflammatory cytokines, and growth factors in the bladder wall were measured by real-time PCR and ELISA. The contractions of bladder strips in response to KCl and carbachol were monitored in OLETF rats.

**Results:** The bladder blood flow was impaired and there was greater ATP release and vesicular nucleotide transporter (VNUT) expression in the OLETF rats than in the LETO rats, but these effects were suppressed by TA administration. Furthermore, the high expression of HIF-1α, 8-OHdG, IL-6, TNF-α, IGF-1, and bFGF in the OLETF rats was reduced by TA administration. In the OLETF rats, the contractile responses of bladder strips to KCl and carbachol were impaired, but were restored by TA administration.

**Conclusions:** The impairment of bladder blood flow in rats with T2DM is associated with greater ATP release and the upregulation of VNUT, markers of hypoxia, proinflammatory cytokines, and growth factors in the bladder epithelium. PDE5 inhibition has the potential to prevent the storage and voiding dysfunction associated with T2DM.

## INTRODUCTION

Many epidemiologic studies have shown that non-urologic disorders, such as hypertension, type 2 diabetes mellitus (T2DM), and dyslipidemia, are associated with lower urinary tract symptoms (LUTS) in both men and women [1]. Lifestyle factors, and especially the presence of T2DM, can affect lower urinary tract function. Such dysfunction has been evaluated by urodynamic studies, which have shown detrusor overactivity (DO) in 55%–61% of individuals with T2DM, detrusor underactivity (DU) in 9%–23%, and areflexia in 9%–23% [2–4]. Furthermore, both DO and DU have been shown to exist in women with T2DM [5]. According to a review by Daneshgari *et al*., a range of lower urinary tract disorders can be present in T2DM, which comprise storage disorders in the early stages and voiding disorders in the later stages [6].

The results of both clinical and basic studies have suggested that atherosclerosis in both men and women induces a decrease in the blood flow in the bladder, leading to chronic ischemia [7]. Chronic bladder ischemia causes oxidative stress, resulting in denervation and the two in the bladder wall, and the progression of DO to DU. However, the mechanisms whereby chronic bladder ischemia causes DO have not been elucidated. In our earlier study of salt-sensitive hypertensive rats, we found that hypertension-related bladder ischemia reduces the bladder capacity of the rats and increases the release of ATP from the bladder epithelium [8]. This provoked the following question: does the same mechanism underlie the development of DO in other models of T2DM?

Most of the research performed using animal models of T2DM has been conducted using rabbits and rats in which arteriosclerosis was artificially induced, but few studies have focused on spontaneous models of T2DM. We used Otsuka Long–Evans Tokushima Fatty (OLETF) rats as a pathologic model of T2DM and Long-Evans Tokushima Otsuka (LETO) rats as controls in our efforts to (i) elucidate the mechanisms of the development of DO and DU in T2DM, and (ii) determine whether DO or DU could be ameliorated by long-term treatment with tadalafil, an inhibitor of phosphodiesterase-5 (PDE5). The OLETF rat model that we used develops insulin resistance by 12 weeks of age and hyperinsulinemia by 25 weeks of age, and subsequently shows a decrease in circulating insulin concentration [9]. In addition, hyperglycemia is maintained throughout this progression. We have published the plasma insulin and glucose concentrations of 36- and 48-week-old OLETF and LETO rats previously [10]. The features of this pathologic model closely resemble those of human T2DM.

Using this rat model of T2DM, we have previously shown that the chronic ischemia caused by T2DM leads to oxidative stress, resulting in the upregulation of several cytokines and growth factors, and prostate enlargement. Furthermore, treatment with a PDE5i ameliorates the prostatic ischemia and might prevent enlargement through the suppression of cytokines and growth factors [11]. Chronic inflammation caused by bladder ischemia may also be associated with the development of overactive bladder (OAB) [12]. Pro-inflammatory cytokines induce the formation of a local “vicious circle,” in which the cytokine interleukin (IL)-8 promotes bronchial smooth muscle contraction and exacerbates asthma symptoms [13], and in addition, in inflammatory bowel disease, IL-17 causes the hypercontraction of small intestinal smooth muscle [14]. Bladder ischemia results in an increase in pro-inflammatory cytokines, but it is not known how this affects bladder function. Therefore, in the present study, we also characterized the proinflammatory cytokine expression in the bladders of rats with T2DM.

## MATERIALS AND METHODS

### Animals

Four-week-old male OLETF and LETO rats were purchased from SLC Inc. (Shizuoka, Japan). The LETO rat is a control strain that does not show features of T2DM. The animals were housed at the University of Fukui Animal Center, at a constant temperature of 23°C and 50%–60% humidity, and under a normal 12-hr light/dark cycle. Tadalafil (100 μg/kg/day) was administered orally to OLETF and LETO rats for 12 consecutive weeks, beginning when they were 36 weeks old, and a control group was administered only vehicle for 12 weeks. Tap water and standard rat chow were provided *ad libitum*. All the animal experiments were conducted in accordance with the guidelines of the Fukui University Committee for Animal Experimentation (Approval number: R03061).

A total of 36 rats were divided into six groups (six rats per group), as depicted in Figure 1A: the L-36 (36-week-old LETO rats), L-48 (48-week-old LETO rats administered vehicle for 12 weeks), L-48(t) (48-week-old LETO rats administered tadalafil for 12 weeks), O-36 (36-week-old OLETF rats), O-48 (48-week-old OLETF rats administered vehicle for 12 weeks), and O-48(t) (48-week-old OLETF rats administered tadalafil for 12 weeks) groups.

**Fig. 1.**
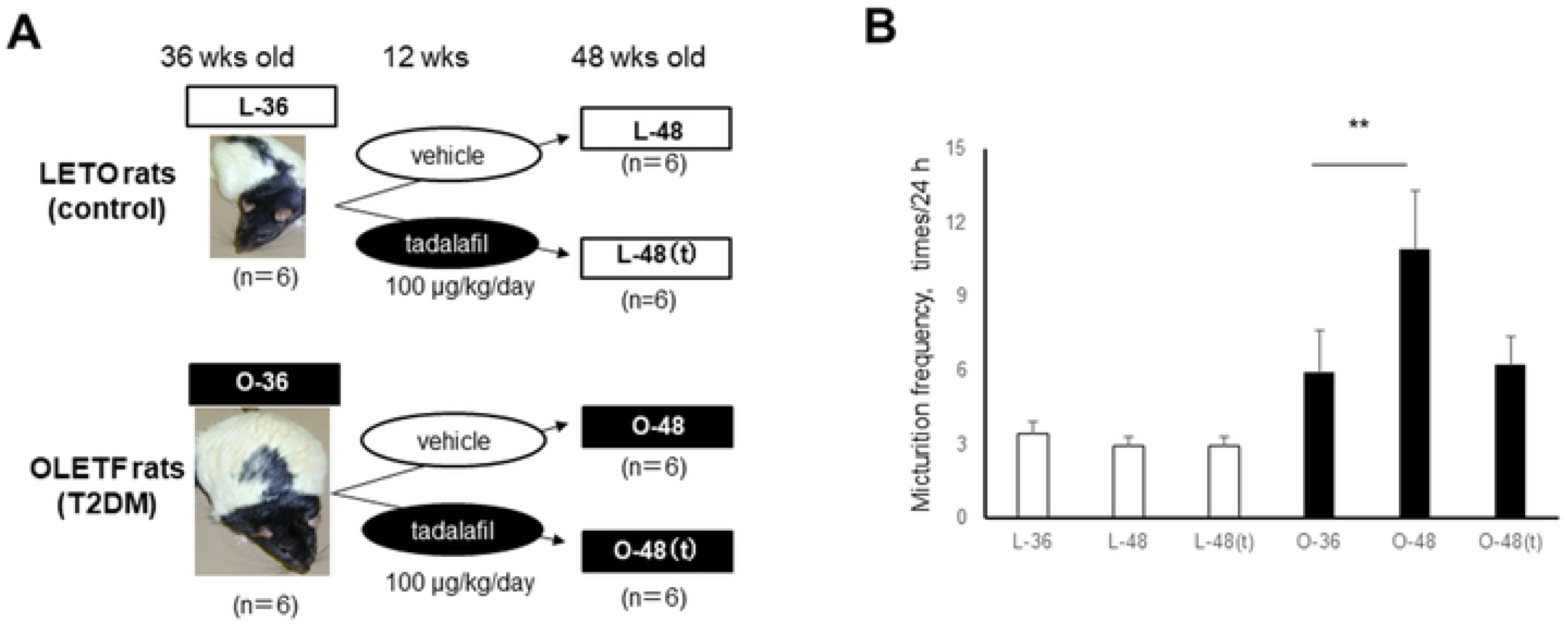
Assessment of the micturition behavior of the rats. **A:** Scheme of the study design, in which OLETF and LETO rats were administered vehicle or tadalafil. L-36: 36-wk-old LETO rats, L-48: 48-wk-old LETO rats administered vehicle for 12 weeks, L-48(t): 48-wk-old LETO rats administered tadalafil for 12 weeks, O-36: 36-wk-old OLETF rats, O-48: 48-wk-old OLETF rats administered vehicle for 12 weeks, O-48(t): 48-wk-old OLETF rats administered tadalafil for 12 weeks. n=6/group. **B:** Frequency of micturition over 24 hr in the LETO and OLETF rats. Data are mean ± SEM. \**p*<0.05, \*\**p*<0.01.

### Micturition behavior

A metabolic cage was used to assess voiding by the rats. The urine voided by each was collected through a urine collection funnel and weighed on an electronic balance, and the cumulative mass of the collected urine was recorded every 10 min. The rats were kept in the metabolic cages for approximately 60 hr to permit acclimation, and the values recorded during the last 24 hr of this period were analyzed. The monitoring period started at 18:00. The data collected were used to calculate the mean volume of urine voided and the number of micturitions per 24-hr period.

### Laser speckle blood flow imaging

The blood flow in the bladders of the LETO and OLETF rats was evaluated using a laser speckle blood flow imaging system, the Omegazone™ (OZ-2, Omegawave Inc., Tokyo), as described by Forrester *et al*. [15]. The rats were anesthetized using halothane, then the bladder was exposed and a catheter was inserted through the bladder dome. The catheter was then connected to an infusion pump, and saline was infused until the intravesicular pressure rose to 10 cm H_2_O. The bladder surface was then diffusely irradiated with a 780-nm semiconductor laser, and the scattered light was passed through a hybrid filter and detected using a charge-coupled device. A single image of the blood flow was generated using the mean of the values obtained from 20 consecutive raw speckle images. The total blood flow to the entire bladder was calculated by adding the values for both components of the blood flow to the bladder.

### Organ bath bladder distention test

After 48-week-old rats were sacrificed by decapitation, their bladders and urethras were removed and weighed. The amount of ATP released from the stretched bladder epithelium was measured using the method of Tanaka *et al*. [16], with slight modifications. One end of the infusion catheter was inserted through the urethra into the bladder and secured at the bladder neck using surgical sutures. After the bladder epithelium had been rinsed three times with 0.5 mL of Krebs solution, the catheter was connected to the infusion pump and pressure transducer. The bladder was then fixed vertically in a 10-mL organ bath containing Krebs solution that was gassed with 5% CO_2_ and 95% O_2_ at 37°C, and physiological saline was then infused into the bladder at a rate of 0.02 mL/min. The bladder was stretched at a pressure of 20 cmH_2_O, which was maintained for 10 min. The solution in the bladder was then collected by gravity and kept on ice. The ATP concentration of this fluid was measured using the ATPlite™ luciferin-luciferase assay with a Fusion luminometer (Perkin Elmer, Waltham, MA, USA), according to the manufacturer’s instructions. The amount of ATP released was then converted to a concentration per mg tissue.

### Real-time PCR

The mRNA expression of genes of interest in the bladders of OLETF rats (O-36, O-48, and O-48(t) groups) was measured using real-time PCR. After the rats had been sacrificed by decapitation, their bladder samples were cut into small pieces and ground into powder using a mortar and pestle under liquid nitrogen. RNA was then isolated from the samples using a RNeasy Fibrous Tissue Mini Kit (Qiagen, Hilden, Germany), according to the manufacturer’s instructions. Two μg of each RNA sample was used for single-strand complementary DNA synthesis using a High-capacity RNA-to-cDNA Kit (Applied Biosystems, Foster City, CA, USA), according to the manufacturer’s protocol, in a final volume of 20 μL. Reverse transcription was performed at 37°C for 60 min, which was followed by inactivation at 95°C for 5 min. The mRNA expression of target genes was quantitatively analyzed using the SYBR green fluorescence method on an ABI 7300 Real-Time PCR System (Applied Biosystems), with GAPDH as the reference gene. The mRNA expression of vesicular nucleotide transporter (VNUT), hypoxia-inducible factor-1 alpha (HIF-1α), interleukin-6 (IL-6), tumor necrosis factor alpha (TNF-α), insulin-like growth factor-1 (IGF-1), and basic fibroblast growth factor (bFGF) in the bladder of each rat was determined, and the fold differences between the groups were calculated. The primers used are listed in Supplemental Table S1.

### ELISA

Bladder tissue samples (15 mg) were cut into small pieces and ground into powder using a mortar and pestle under liquid nitrogen. The soluble and insoluble protein fractions were extracted using a CelLytic™ MEM Protein Extraction Kit (Sigma–Aldrich, St. Louis, MO, USA) and a CelLytic™ NuCLEAR™ Extraction Kit (Sigma-Aldrich), respectively, according to the manufacturer’s instructions. The protein levels of IL-6, TNF-α, IGF-1, and bFGF were measured using a Rat IGF-1 ELISA Kit (ab213902, Abcam, Cambridge, UK), a Rat IL-6 ELISA Kit (ab100772, Abcam), a Rat TNF alpha ELISA Kit (ab100785, Abcam), a Rat bFGF ELISA Kit (Invitrogen, CA), and an AssayMax™ Dihydrotestosterone ELISA Kit (Assaypro, St. Charles, MO, USA), respectively. The protein concentration of each was calculated as picograms of protein per milligram of tissue wet mass.

### Measurement of 8-hydroxy-*2’*-deoxyguanosine (8-OHdG) concentration

To measure the 8-OHdG concentration, bladder tissue samples were homogenized in 0.1 M phosphate buffer containing 1 mM EDTA. After centrifugation for 10 min, the supernatant was removed and purified using a DNeasy Blood & Tissue Kit (Qiagen), according to the manufacturer’s instructions, then the DNA was digested using nuclease P1 (Wako, Tokyo). After the addition of 1 unit of alkaline phosphatase (Wako) per 100 μg of DNA and incubation at 37°C for 30 min, the samples were boiled for 10 min and kept on ice until use. The concentration of 8-OHdG was then measured using an 8-Hydroxy-2-deoxyguanosine ELISA kit (ab201734, Abcam).

### In vitro bladder-strip experiments

For the *in vitro* bladder-strip experiments, rats in the O-36, O-48, and O-48(t) groups were used that were different to those used for the measurement of ATP release from the bladder epithelium. After the rats had been sacrificed by decapitation, full-thickness longitudinal bladder strips (7 mm long × 3 mm wide) were excised and mounted in a 10-mL organ bath containing Krebs solution at 37°C that was continuously gassed with 95% O_2_ and 5% CO_2_. The strips were equilibrated under a resting tension of 1 g for 45 min, with the Krebs solution being changed every 15 min. Following equilibration, the strips were exposed to a solution containing a high K^+^ concentration (62 mM KCl) to normalize the contractile force. After washing away the KCl solution, the isometric contractions of the strips caused by the cumulative application of carbachol (3×10^−7^, 10^−6^, 3×10^−6^, 10^−5^, and 3×10^−5^ M) or atropine (0, 10^−7^, 10^−9^, 10^−8^, and 10^−7^ M) were recorded *via* force transducers (Nihon Kohden, Tokyo, Japan) and Labchart & Scope software (ADInstruments, Colorado Springs, CO, USA). At the end of this experiment, the bladder strips were weighed, and the contractile force of each was normalized to its mass.

### Data analysis

The results are presented as the mean ± standard error of the mean (SEM). All of the data were analyzed using analysis of variance (ANOVA) and the independent-samples *t*-test using SPSS^®^ ver. 16.0J for Windows^®^ (SPSS, Inc., Chicago, IL, USA). *P* < 0.05 was regarded as indicating statistical significance.

## RESULTS

### Body masses, bladder wet masses, and micturition characteristics of LETO and OLETF rats

The body masses of the LETO rats and OLETF rats were 546 ± 28 g and 671 ± 31 g (*p*<0.01) at 36 weeks of age and 557 ± 27 g and 658 ± 32 g (*p*<0.01) at 48 weeks of age, respectively. Thus, those of the OLETF rats were significantly higher than those of the LETO rats. The respective wet masses of the bladders of the LETO rats and OLETF rats were 86 ± 3 mg and 175 ± 7 mg (*p*<0.01) at 36 weeks of age and 103 ± 5 mg and 196 ± 14 mg at 48 weeks of age (*p*<0.01), respectively. Those of the OLETF rats were thus also significantly heavier. Tadalafil administration did not affect the body masses or bladder wet masses of either group. The mean voided urine volume did not significantly differ between the LETO and OLETF rats, and this was also not affected by age or tadalafil administration.

The frequency of micturition during a 24-hr period of the OLETF rats was significantly higher than that of the LETO rats at both 36 and 48 weeks of age, and the frequency of micturition of the OLETF rats was significantly higher at 48 weeks of age than it was at 36 weeks of age (*p*<0.01). Tadalafil administration significantly reduced the frequency of micturition of the 48-week-old OLETF rats (*p*<0.05), but the frequency of micturition of the LETO rats did not significantly differ between 36 and 48 weeks of age and was not affected by tadalafil administration (Fig. 1B).

### ATP production and VNUT expression in the bladder

To clarify the mechanisms underlying the improvement in the urinary frequency of the OLETF rats induced by tadalafil, we measured the amounts of ATP released from the bladder epithelium by means of an organ bath bladder distention test (Fig. 2A). The ATP concentrations of the OLETF rats were significantly higher than those of the LETO rats (*p*<0.01), and the concentrations in both the LETO rats (*p*<0.05) and OLETF rats (*p*<0.01) were significantly reduced by tadalafil (Fig. 2B). Thus, the tadalafil-induced reduction in urinary frequency of the OLETF rats was accompanied by a reduction in ATP release from the bladder epithelium. To investigate the mechanisms underpinning the changes in ATP production, we measured the expression of VNUT in the bladder. At 36 weeks of age, the VNUT expression of the OLETF rats was significantly higher than that of the LETO rats (*p*<0.05) (Fig. 2B), but by 48 weeks of age, the VNUT expression of the LETO rats had increased (*p*<0.05), as had that of the OLETF rats (*p*<0.01). Moreover, tadalafil treatment significantly reduced the VNUT expression in both types of rats (*p*<0.01). Thus, the change in bladder VNUT expression might underpin the changes in the production of ATP in the bladder epithelium.

**Fig. 2.**
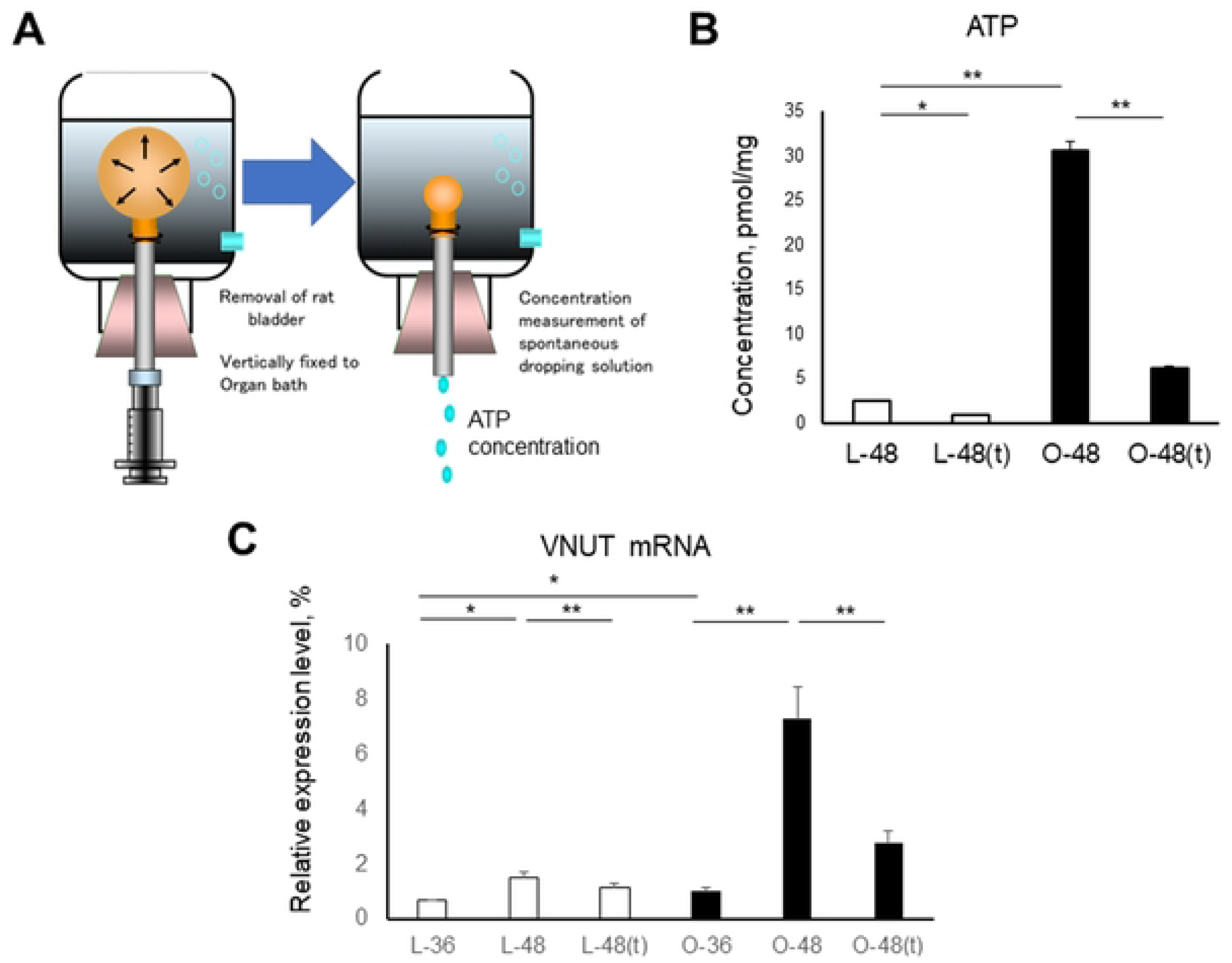
Amount of ATP released in response to bladder distention and the expression of VNUT in the bladder. **A:** Measurement of distention in an organ bath. A whole bladder was placed in an organ bath and injected with saline, which was collected by gravity after the syringe was removed. **B:** Amount of ATP released from the stretched bladder epithelium of 48-week-old LETO and OLETF rats during distention at 20 cm H_2_O pressure for 10 min. **C:** VNUT mRNA expression in the bladder in 36- and 48-week-old LETO rats. Data are mean ± SEM. \**p*<0.05, \*\**p*<0.01.

**Fig. 3.**
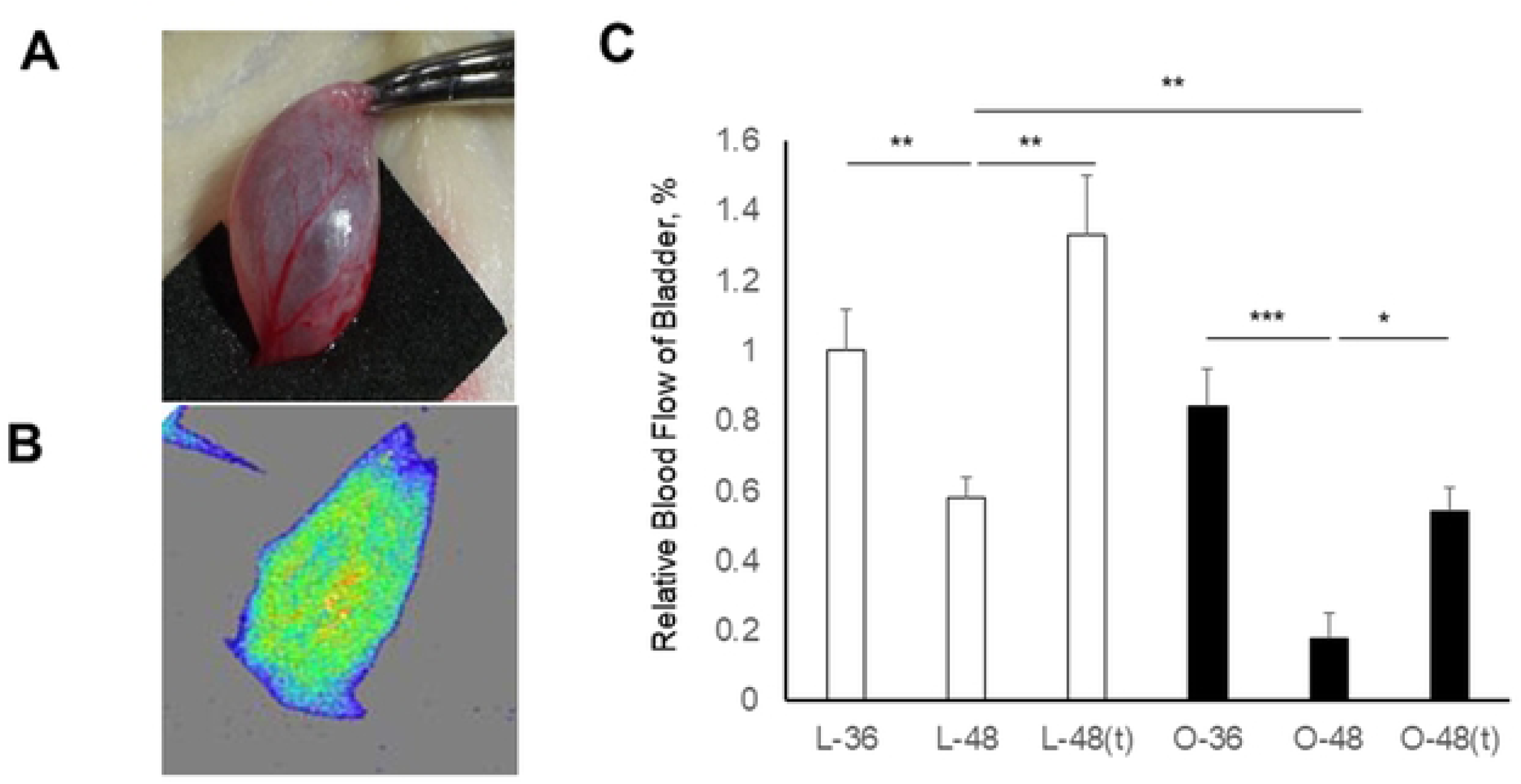
Blood flow to the bladder in LETO and OLETF rats. **A:** Image of a rat bladder dissected and exposed under anesthesia. **B:** Laser speckle image of a rat bladder. **C:** Comparison of the bladder blood flow in the six groups of rats. The blood flow is expressed as a percentage relative to that of the L-36 rats. n=6/group. Data are mean ± SEM. \**p*<0.05, \*\**p*<0.01.

### Changes in the blood flow to the bladder, assessed using laser speckle blood flow imaging

The blood flow in the bladder was assessed using the Omegazone laser speckle blood flow imaging system just before the bladder was extracted from each of the rats (Fig 2A). The blood flow was displayed as changes in the laser frequency in the speckle imaging. The Omegazone uses pixels of different colors, with blue pixels indicating areas with poor blood flow and yellow pixels indicating those with good blood flow (Fig 2B). The imaging revealed that the blood flow to the bladder was significantly lower in the 48-week-old LETO rats than in 36-week-old rats (*p*<0.01), and also in the OLETF rats (*p*<0.005). At 48 weeks of age, the blood flow in the OLETF rats was significantly lower than that in the LETO rats (*p*<0.01). Tadalafil administration caused significant increases in the blood flow to the bladder in both the LETO (*p*<0.01) and OLETF (*p*<0.05) rats (Fig. 2C). These results indicate that the blood flow to the bladder of older OLETF rats is lower than that of LETO rats, but it is increased by tadalafil.

### Expression of HIF-1α and the bladder 8-OHdG concentration

To evaluate the hypoxia of the bladder induced by the changes in blood flow, we evaluated the expression of HIF-1α in the bladders of OLETF rats, and found that it was significantly higher in 48-week-old rats than in 36-week-old rats (*p*<0.01). In addition, HIF-1α expression tended to be reduced by tadalafil, but this effect was not statistically significant (Fig. 4A). 8-OHdG is a marker of oxidative stress that is produced when DNA is damaged by reactive oxygen species. The 8-OHdG concentrations in the bladders of OLETF rats were significantly higher in 48-week-old rats than in 36-week-old rats (*p*<0.01), and were significantly reduced by tadalafil (*p*<0.01) (Fig. 4B). These results indicate that hypoxia is induced by the reduction in the blood flow to the bladders of older OLETF rats.

**Fig. 4.**
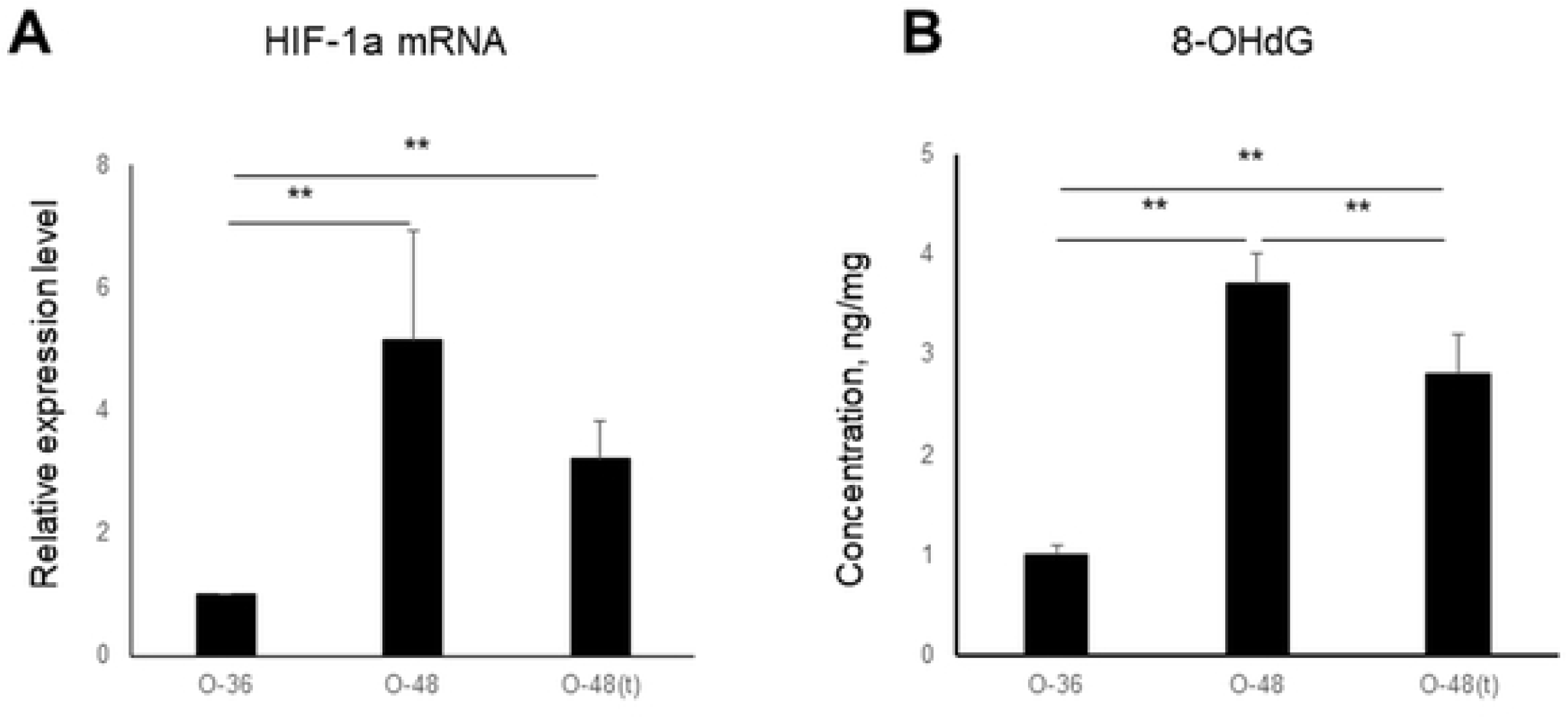
HIF-1α mRNA expression **(A)** and 8-OHdG concentration **(B)** in the bladder of OLETF rats at 36 weeks and 48 weeks of age, in the presence and absence of tadalafil. n=6/group, and two replicates were obtained per sample. Data are mean ± SEM. \**p*<0.05, \*\**p*<0.01.

### mRNA and protein expression of pro-inflammatory cytokines and growth factors in the bladder

The expression of the proinflammatory cytokines IL-6 and TNF-α and the growth factors IGF-1 and bFGF, which are associated with bladder inflammation, were next measured in OLETF rats. The mRNA expressions of IL-6, TNF-α, IGF-1, and bFGF were significantly higher in 48-week-old rats than in 36-week-old rats (*p*<0.01), and they tended to be reduced by tadalafil. The effect of tadalafil was significant for TNF-α expression (*p*<0.01) (Fig. 5A). The protein expressions of IL-6, TNF-α, IGF-1, and bFGF were also significantly higher in 48-week-old rats than in 36-week-old rats (*p*<0.01), and tended to be reduced by tadalafil. The effects of tadalafil were statistically significant for IL-6, TNF-α, and IGF-1 (*p*<0.05 or *p*<0.01). Thus, the expression of proinflammatory cytokines and growth factors changes with age in OLETF rats and is affected by tadalafil.

**Fig. 5.**
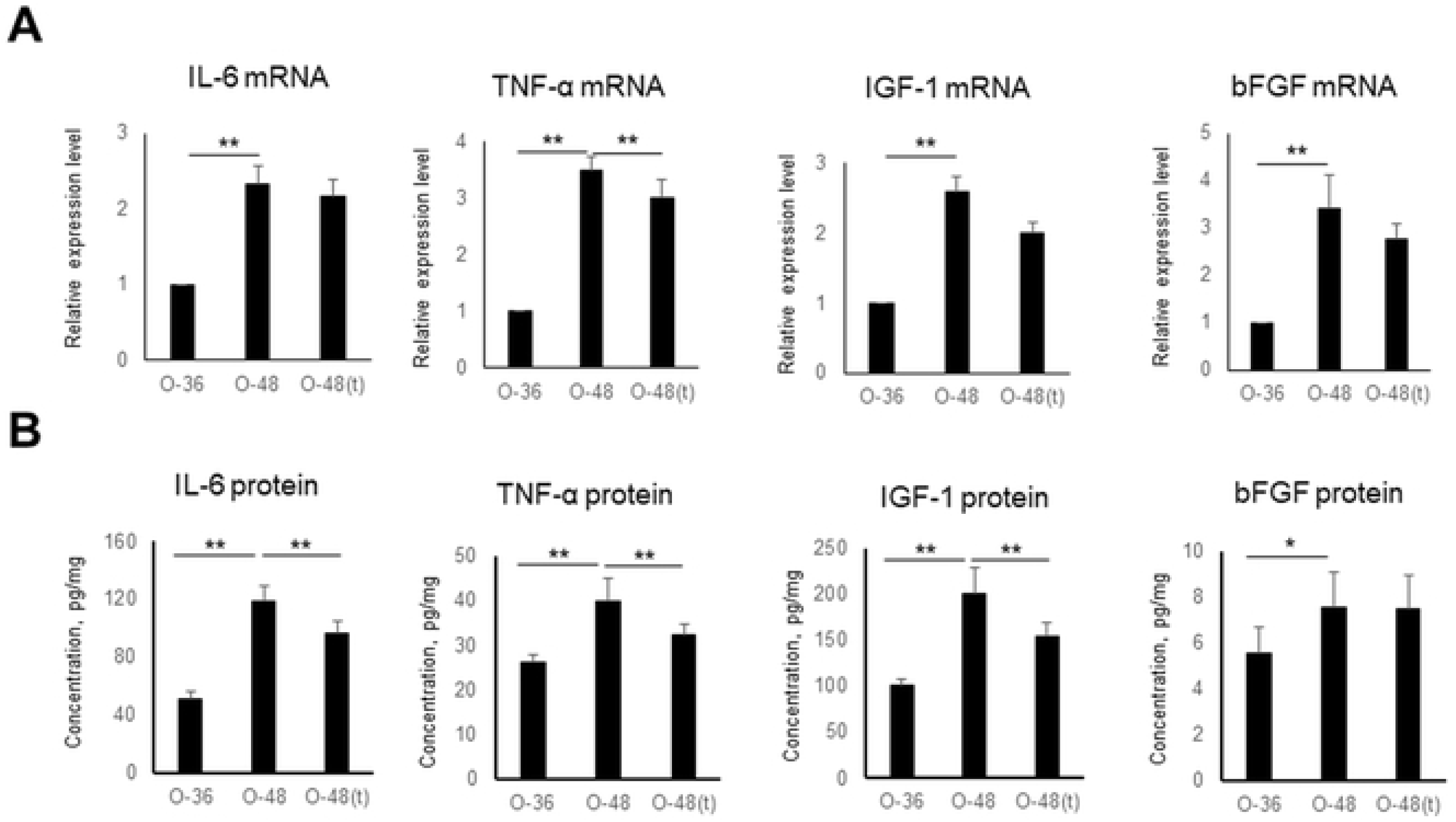
mRNA expression, normalized to that of *Gapdh* **(A)** and protein concentrations **(B)** of proinflammatory cytokines (IL-6 and TNF-α) and growth factors (IGF-1 and bFGF) in the bladder of OLETF rats at 36 and 48 weeks of age, in the presence or absence of tadalafil, measured using real-time PCR and ELISA, respectively. n=6/group, and two replicates were obtained per sample. Data are mean ± SEM. \**p*<0.05, \*\**p*<0.01.

### Contractile responses of bladder strips to KCl, carbachol, and atropine

We used bladder strips obtained from OLETF rats to evaluate the bladder contraction responses to KCl and carbachol, and found that the response to KCl was significantly weaker in 48-week-old rats than in 36-week-old rats (*p*<0.05). In addition, tadalafil significantly increased the bladder contractions (*p*<0.01) (Fig. 6A). We also found that the bladder showed a dose-dependent response to carbachol, but the sensitivity of the bladder to carbachol was lower in rats of 48 weeks of age than in those of 36 weeks of age. However, the sensitivity of the bladder in rats of 48 weeks of age was restored by tadalafil (Fig. 6B). The response of the bladder to KCl was reduced in a dose-responsive manner by atropine administration, but the sensitivity of the bladder to atropine was lower in 48-week-old rats than in 36-week-old rats. However, the sensitivity of the bladders of 48-week-old rats was restored by tadalafil (Fig. 6C). Thus, the contractile response of the bladders of OLETF rats decreases in old age, but is restored by tadalafil.

**Fig. 6.**
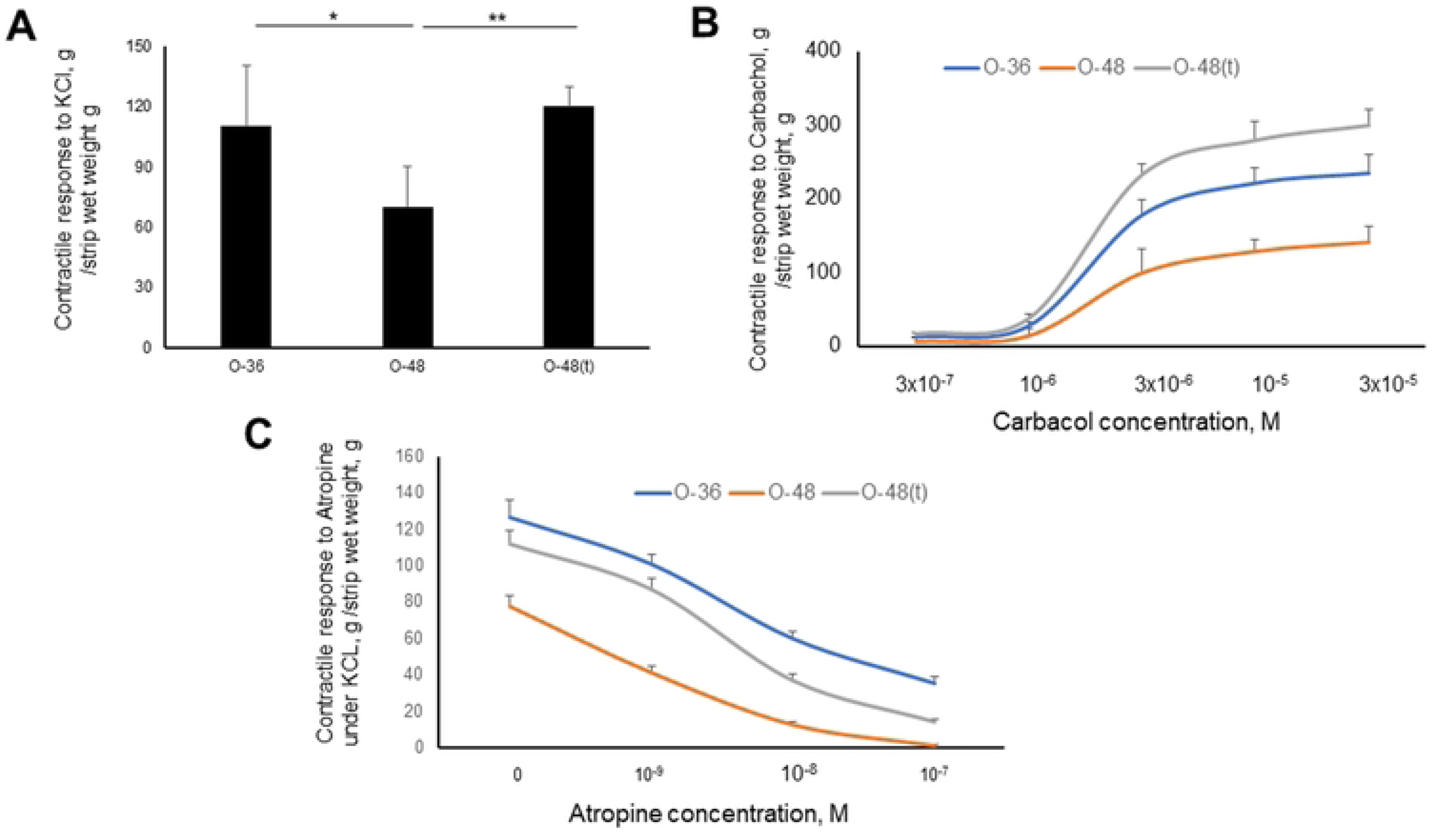
Contractile responses of bladder strips to 62 mM KCl **(A)**, carbachol (3×10^−7^, 10^−6^, 3×10^−6^, 10^−5^, 3×10^−5^M), **(B)** and atropine (0, 10^−7^, 10^−9^, 10^−8^, 10^−7^ M) in the presence of 62 mM KCL **(C)**. The strips were obtained from OLETF rats at 36 and 48 weeks of age, in the presence or absence of tadalafil. n=6/group, and two replicates were obtained per sample. Data are mean ± SEM. \**p*<0.05, \*\**p*<0.01.

## DISCUSSION

In the present study, we have shown that in a rat model of T2DM, the expression of HIF-1α and various proinflammatory cytokines and growth factors, and the concentration of 8-OHdG are increased as a result of abnormal bladder perfusion. The increase in the production of proinflammatory cytokines may increase the release of ATP from the epithelium of the bladder *via* the upregulation of VNUT. In addition, we found that the contractile responses of the bladder to KCl and muscarinic stimulation decreased with age. Long-term (12-wk) administration of tadalafil improves the blood flow to, and ameliorates the oxidative stress in, the bladder, resulting in the release of less ATP from the epithelium and an improvement in bladder responsiveness. This study is the first to provide evidence that epithelium-derived ATP may be associated with abnormal urine storage in T2DM and that tadalafil inhibits ATP release and improves bladder responsiveness.

Diabetes can cause lower urinary tract dysfunction, including DO or DU [6,17]. Many studies have investigated the bladder function and morphology of animal with streptozotocin-induced type 1 diabetes mellitus (T1DM), most of which are characterized by the presence of bladder hypertrophy [18]. Two genetic models of T2DM, Zucker Diabetic Fatty rats and *db*/*db* mice, which have blood glucose concentrations that are similar to those of the T1DM model, show the same degree of, or only slight, hypertrophy [19]. In contrast, the Goto-Kakizaki rat, a genetic model of T2DM, shows mild-to-moderate bladder hypertrophy. In the present study, the model of T2DM (OLETF rats) that was used showed an increase in bladder wet mass with age and bladder hypertrophy *vs*. control LETO rats. Moreover, the daily urine production by the OLETF rats tended to be larger than that by the LETO rats. Diabetic polyuria induces DNA synthesis in the bladder, resulting in greater protein synthesis and hyperplasia of the smooth muscle and epithelial layers of the wall [20].

Although OAB is present from an early stage of T2DM in humans, the presence of DO has not been reported following the cystometric evaluation of an animal model of T2DM [20]. In rats fed a fructose-rich diet, no abnormality in bladder capacity has been identified, but more frequent non-voiding contractions during the storage phase have been identified cystometrically [21]. Nevertheless, quantitative data regarding non-voiding contractions have not been generated [22]. In the present study, we found that the mean volume voided by LETO and OLETF rats did not differ and non-voiding contractions were not observed prior to the activation of the micturition reflex during cystometric evaluation.

The bladder epithelium plays an important role in mechanosensory transduction [23,24]. The production of ATP from epithelial cells, which is induced by mechanical stimuli, activates purinergic receptors on submucosal afferent fibers, facilitating bladder sensory signaling [25]. There is substantial release of ATP from the bladder epithelium in the presence of various pathologic conditions in humans, such as OAB, benign prostatic hyperplasia, spinal cord injury, and interstitial cystitis [26–28]. We have previously demonstrated that epithelium-derived ATP is more abundant in salt-sensitive hypertensive rats, and that this is accompanied by lower mean voided volume [8]. The present study is the first to demonstrate greater ATP release from the bladder epithelium in an animal model of T2DM. However, the reason for this greater ATP release in T2DM remains to be determined. Because PDE5 inhibition improved the blood flow to the bladders of the rats and reduced the release of ATP, it can be speculated that, at least in this model, hypoxia triggers a release of ATP from the epithelium. Nephropathy, retinopathy, and neuropathy, which are the three major complications of T2DM, develop when vascular endothelial cells are exposed to hyperglycemia, which impairs the microcirculation. This condition can progress to cause macrovascular complications, including coronary artery and cerebrovascular abnormalities [29]. The rat model of T2DM used in the present study exhibited bladder ischemia, which was associated with higher expression of HIF-1α and various proinflammatory cytokines and growth factors, and higher 8-OHdG concentrations in the bladder wall. We speculate that diabetes increases the ATP release from the epithelium through the mechanism described below.

Several mechanisms have been postulated to explain the release of ATP from the bladder epithelium during bladder distention, including vesicular exocytosis and the provision of connexin/pannexin channels [30–33]. It has been suggested that ATP release from its storage vesicles is regulated principally by VNUT (gene name: *Slc17A9*) in the bladder epithelium [34]. VNUT mediates the active transport of ATP that concentrates it in its secretory vesicles [36,36]. Slight stretching of bladder epithelium harvested from VNUT knockout mice reduces the release of ATP, suggesting that VNUT-mediated ATP release is involved in the mechanism of the bladder relaxation that occurs during the early stages of filling [34]. In contrast, ATP release in response to lipopolysaccharide is incompletely blocked by the inhibition of pannexin channels [31,33], but the ATP release evoked by bladder distention is completely inhibited by antagonists and by the small interfering RNA-mediated knockdown of pannexin [37]. The results of these studies suggest that there are multiple mechanisms for ATP release from the bladder epithelium that may be activated by differing stimuli. In the present study, the ATP release evoked by bladder distention was more substantial in the OLETF rats than in the LETO rats. In addition, the expression of VNUT was much higher in the bladders of the OLETF rats, which implies that the mechanism of ATP release may shift from transport through connexin/pannexin channels to vesicular exocytosis. However, the trigger for this transition has not been firmly established

A few previous studies have shown that ATP release is stimulated by inflammation. Mice with conditional hepatic double-knockout of their *Irs1* and *Irs2* genes showed characteristics typical of T2DM and OAB [38]. In this model, when the hyperactivated TNF-α-mediated signaling in the bladder of the mice was inhibited, the OAB was ameliorated, without any effect on the hyperglycemia. In the present study, the expression of IL-6, TNF-α, IGF-1, and bFGF in the bladders of the OLETF rats increased with age, but was reduced by tadalafil administration. However, further studies are necessary to elucidate the relationship between inflammation and ATP release from the bladder epithelium.

Although the nitric oxide (NO)/cyclic guanosine monophosphate (GMP) pathway is an endothelium-derived signal that induces vascular smooth muscle relaxation, the results of two previous studies suggest that it is also involved in bladder relaxation [39,40]. Another PDE5 inhibitor, sildenafil, has been shown to increase the cGMP concentration and inhibit the distention-induced release of ATP from the bladder epithelium in mice [39], and to significantly reduce the amount of ATP released in response to lipopolysaccharide administration. Furthermore, an earlier study showed that sildenafil inhibits the release of ATP from mucosa isolated from the detrusor muscle [41]. These findings suggest that the NO-cGMP pathway may reduce epithelial ATP release.

A possible mechanism by which PDE5 inhibition reduces the release of ATP is the attenuation of the Ca^2+^ influx *via* transient receptor potential vanilloid (TRPV) 2 and 4 [40]. In the present study, 12 weeks of tadalafil administration markedly reduced ATP release from the bladder epithelium, but it is unclear whether this was an acute effect, exerted *via* the NO-cGMP pathway, or a chronic effect, associated with lower proinflammatory cytokine production, secondary to an improvement in blood flow. Because the effect occurred >2 days after the completion of the tadalafil administration, and because the interiors of the bladders were rinsed three times with 0.5 mL of Krebs solution, we speculate that the effect may be the result of an improvement in blood flow to the bladder.

We also found that bladder strips from 48-week-old rats showed poorer contractile responses to KCl and carbachol than strips from 36-week-old rats, suggesting that hyposensitivity to muscarinic stimulation and/or a decrease in the expression of contractile proteins, alongside an increase in collagen deposition, in bladder smooth muscle occurs during the later stage of T2DM. We did not evaluate the histology of the bladders of the rats, but it is known that the bladder becomes less responsive to muscarinic stimulation with age, and that this is accompanied by bladder hypertrophy. The later stage of diabetic bladder dysfunction has been described as dysfunctional voiding, owing to underactivity of the bladder [6]. This is presumed to be caused by chronic exposure to oxidative stress, secondary to long-term hyperglycemia [42]. In rats, chronic bladder ischemia induced by endothelial injury of the iliac arteries, has been shown to reduce the contractile responses of bladder strips to KCl, electrical stimulation, and carbachol, and to be accompanied by collagen deposition in the smooth muscle layer. In this study, 8 weeks of tadalafil administration prevented the decline in bladder contractility and the collagen deposition. In the present study, we found that tadalafil administration from 36 weeks of age can also prevent the abnormal bladder contraction that is associated with T2DM in rats. Thus, PDE5 inhibitors may be useful treatments for the abnormal voiding associated with T2DM.

### Limitations

In the present study, we conducted an *in vitro* contraction study of the detrusor muscle to assess bladder function, but not a histologic examination of the bladder wall. We cannot say with certainty why the contractions were attenuated in response to carbachol. Thus, an important question remains: why is T2DM associated with greater release of ATP from the bladder epithelium? Although pro-inflammatory cytokines (IL-6 and TNF-α) or growth factors (IGF-1 and bFGF) are candidate mediators, cytokine-specific inhibitors are needed to assess this possibility.

## CONCLUSIONS

We have studied OLETF rats, which recapitulate a series of the pathologic conditions associated with T2DM, including bladder wall ischemia, high circulating proinflammatory cytokine and growth factor concentrations, the release of ATP from the epithelium, and an impairment in bladder contractility. Tadalafil, a PDE5 inhibitor, ameliorated the ischemia, reduced the concentrations of proinflammatory cytokines, inhibited the release of ATP from the bladder epithelium, and improved bladder contractility. This model should be useful in the further elucidation of the pathogenesis of the bladder dysfunction that is associated with T2DM and contribute to the design of new therapeutic strategies.

## ACKNOWLEDGEMENTS

We thank all of the project members. We also thank Mark Cleasby, PhD from Edanz (https://jp.edanz.com/ac) for editing a draft of this manuscript.

## AUTHOR CONTRIBUTIONS

Takafumi Kabuto, So Inamura, Osamu Yokoyama, and Naoki Terada conceived and designed the study. Takefumi Kabuto analyzed the results and wrote the manuscript. Hisato Kobayashi, Xinmin Zha, Keiko Nagase, Minekatsu Taga, Masaya Seki, Nobuki Tanaka, and Yoshinaga Okumura collected data. Osamu Yokoyama critically revised the manuscript. Naoki Terada supervised the project.

