## Supplementary figures and images for "Phosphodiesterase-5 inhibition inhibits epithelial ATP release and restores detrusor contractility in rats with type 2 diabetes *via* an increase in bladder blood flow"

### Supplment table 1

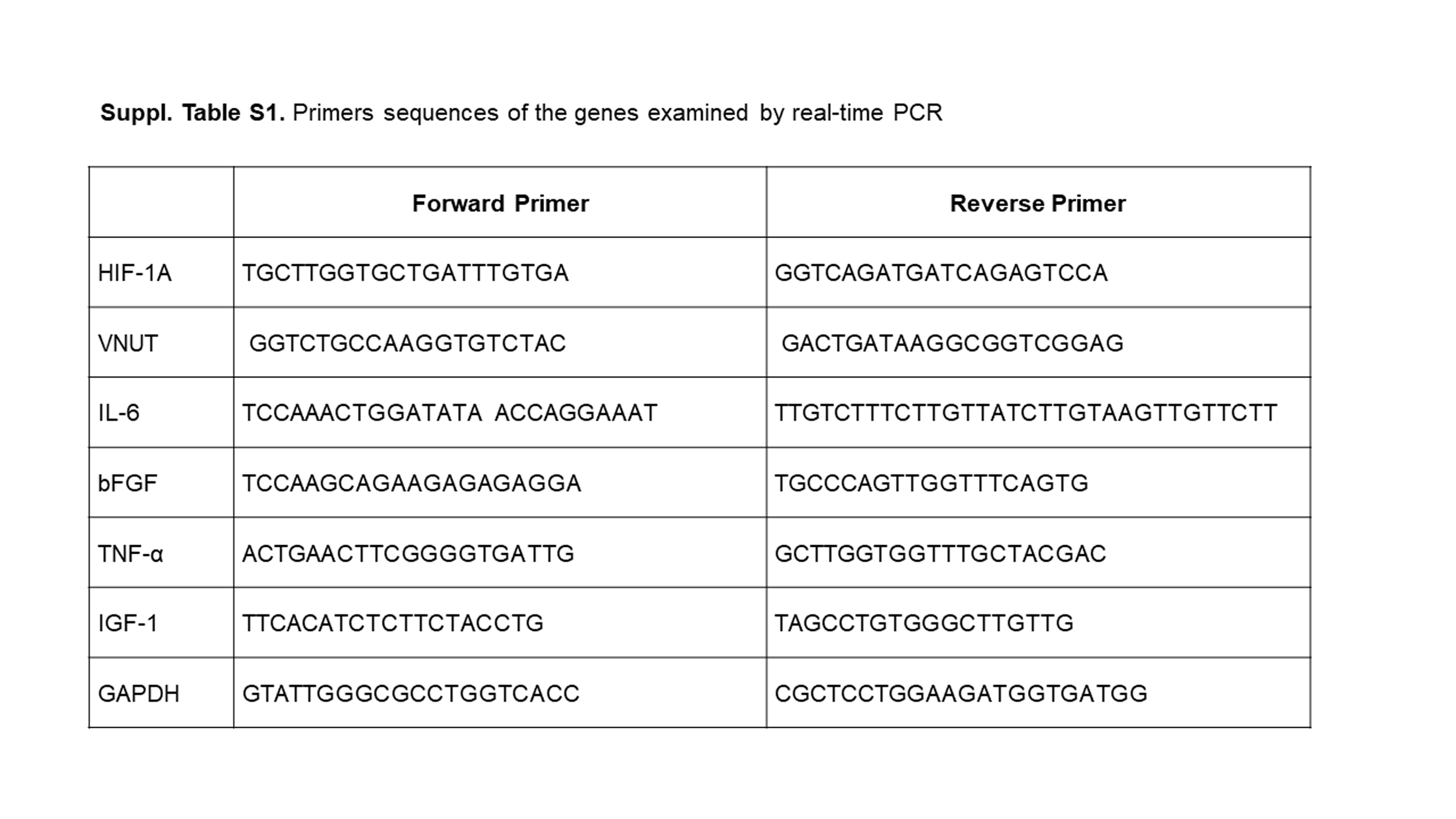
